# *CROSSalive*: a web server for predicting the *in vivo* structure of RNA molecules

**DOI:** 10.1101/626085

**Authors:** Riccardo Delli Ponti, Alexandros Armaos, Gian Gaetano Tartaglia

**Author notes:** Corresponding author: Gian Gaetano Tartaglia.

## Abstract

Here we introduce *CROSSalive*, an algorithm to predict the RNA secondary structure profile (double and single stranded regions) *in vivo* and without sequence length limitations. Using predictions of protein interactions *CROSSalive* predicts the effect of N_6_ adenosine methylation (m6a) on RNA structure. Trained on icSHAPE data in presence (m6a+) and absence (m6a-) of methylation *CROSSalive* achieves an accuracy of 0.88 on the test set. The algorithm was also applied to the murine long non-coding RNA *Xist* (17900 nt, not present in the training) and shows a Pearson’s correlation of 0.45 with SHAPE-map data. *CROSSalive* webserver is freely accessible at the following page: http://service.tartaglialab.com/new_submission/crossalive

## INTRODUCTION

The *in vitro* RNA structure differ from that *in vivo* for the action of interactions such as RNA-binding proteins (Livi *et al.*, 2015). The complex mechanisms contributing to the formation of structure *in vivo* are poorly characterized and previous analysis suggests a prevalence of single stranded regions for all RNA types (Rouskin *et al.*, 2014), although conservation of double-stranded regions has been observed for some non-coding RNAs (Spitale *et al.*, 2015). In the cellular environment RNA undergoes a number of modifications such as methylation that can influence the stability and turnover of the whole transcriptome (Liu and Jia, 2014). *Mettl3* is the key component of the complex that methylates adenosine residues at N_6_ (m6a) and plays a central role in determining RNA structure. Indeed, a method of probing RNA structure using the chemical probe NAI-N3, icSHAPE, indicated that m6a promotes transition from double-to single-stranded regions (Spitale *et al.*, 2015). Through analysis of icSHAPE data, we developed the *CROSSalive* method for the prediction of RNA secondary structure *in vivo*. One key element of our approach is the use of *cat*RAPID predictions of protein interactions to classify single- and double-stranded regions of RNA molecules (Bellucci *et al.*, 2011). *cat*RAPID builds on top of accurate RNA structure calculations performed with the algorithm *RNAfold* (Lorenz *et al.*, 2011).

## METHODS

*CROSSalive* profiles a RNA sequence computing the corresponding secondary structure *in vivo* with (m6a+) and without methylation (m6a-). The algorithm uses predictions of protein interactions to assign single- and double-stranded regions (Spitale *et al.*, 2015):

- For the training and testing we selected RNA fragments carrying the central nucleotide with the highest (single-stranded; 10^5^ sequences) and lowest icSHAPE reactivity (double-stranded; 10^5^ sequences). Each RNA fragment contains 51 nucleotides to allow calculations of protein interactions using *cat*RAPID (Bellucci *et al.*, 2011). The nucleotides are represented as 4-mers: A = (1, 0, 0, 0), C = (0, 1, 0, 0), G = (0, 0, 1, 0) and U = (0, 0, 0, 1).
- The *cat*RAPID approach uses a phenomenological potential that exploits *RNAfold* algorithm to provide accurate information on RNA structure (Bellucci *et al.*, 2011). 7797 regions from a library of 640 canonical RNA-binding proteins (Agostini *et al.*, 2013) were analyzed to identify those able to discriminate nucleotides in single and double-stranded state with an accuracy > 0.6 (m6a+: 228 regions; m6a-: 206 regions).
- The dataset is enriched for proteins with gene ontology (Klus *et al.*, 2015) related to RNA structure (double- and single-stranded RNA binding; helicase activity; m6a+: 101 regions; m6a−: 81 regions; Supplementary Data). The Youden cut-off was computed on the *cat*RAPID scores for each protein in the dataset. Scores above the cut-off were set to 1 (0 otherwise).
- Neural networks (m6a+ and m6a−, with and without protein contributions) were trained using the architecture described in our previous publication for icSHAPE *in vitro* data prediction (Delli Ponti *et al.*, 2017) and cross-validated against each other (Supplementary Data). Each RNA fragment is assigned a score between −1 (high propensity to be single-stranded) to 1 (high propensity to be double-stranded).

## RESULTS

*CROSSalive* scores were ranked by their absolute value and equal groups of positives and negatives were selected to assess the overall performances of the algorithm. From low (50%) to high-confidence (HC) scores (1%, Figure 1A) the accuracy of all the models increases monotonically reaching a maximum of 0.86 for the m6a+ one when protein interactions are used (10-fold cross-validation, CV). In comparison, the *in vitro* icSHAPE model based on RNA sequence information only (Delli Ponti *et al.*, 2017) discriminates single- and double-stranded regions with a 0.88 accuracy (10-fold CV on 1% HC scores). The m6a− *in vivo* model shows lower accuracy (0.74 in 10-fold CV on 1% HC scores) most likely because m6a removal affects the quality of the training set by altering the stability and turnover of the transcriptome (Liu and Jia, 2014). We applied *CROSSalive* to an independent *in vivo* SHAPE-Map experiment (Smola *et al.*, 2016) on the long non-coding *Xist* (17900 nt; not in the training). We used the *in vivo* m6a− model because *Mettl3* is poorly abundant in the trophoblasts (Thul *et al.*, 2017) employed in SHAPE-Map and only few nucleotides are methylated at the 5’ and 3’ of *Xist* (Patil *et al.*, 2016). *CROSSalive* profile shows a correlation of 0.45 with the SHAPE-Map one (Figure 1B). The algorithm achieves an Area under the ROC curve (AUC) of 0.83 on the 15% HC single- and double-stranded regions ranked by SHAPE reactivity. By contrast, the m6a-model trained on RNA sequence information only achieves an AUC of 0.53.

**Figure 1.**
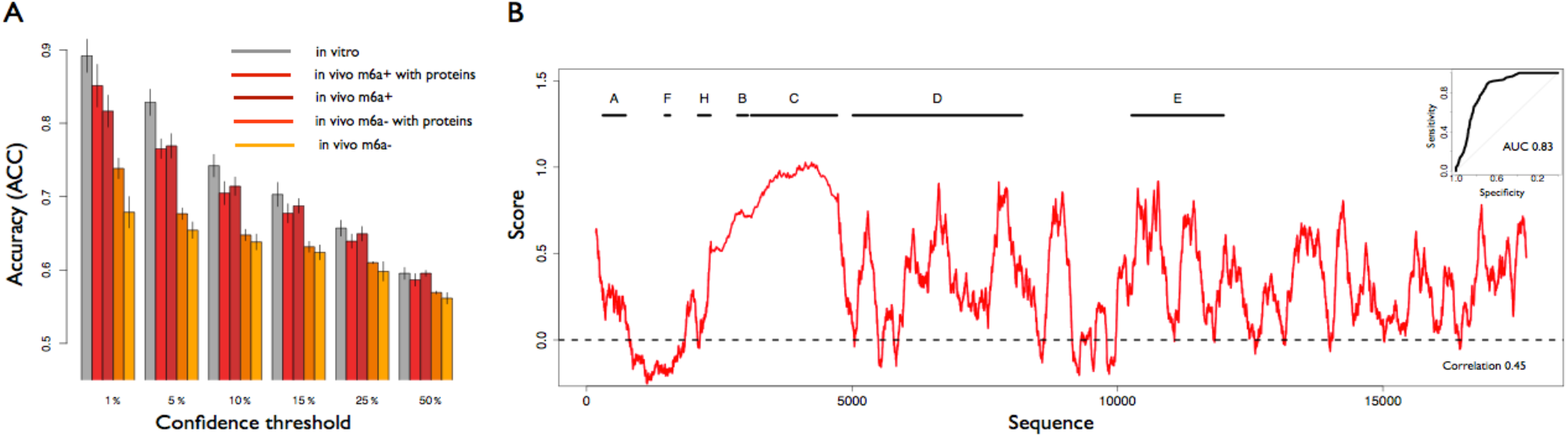
CROSSalive performances. (A) 10-fold cross validation for each specific algorithm (in vitro, in vivo m6a+, in vivo m6a−) with the same training and testing conditions (balanced training set, filtering out sequence redundancy). The accuracies are reported for the scores ranked by their absolute value (same number of positives and negatives were selected), where 50% is the complete set (median). Integrating predictions with protein interactions improves the accuracy. (B) Secondary structure profile of Xist using m6a− model. Known repetitive regions of Xist such as Rep A and Rep C are reported to be very structured (i.e. score > 0). The predicted profile has an overall correlation of 0.45 with in vivo SHAPE data. In the top right we report the ROC curve of CROSSalive on the top and bottom 15% ranked SHAPE data (AUC of 0.83).

## CONCLUSIONS

By using sequence-based information, *CROSSalive* profiles the RNA secondary structure *in vivo*. The use of different models (*in vivo / in vitro*, m6a+ / m6a-) could help to identify structural regions to investigate experimentally. As previously done with *CROSS* (Delli Ponti *et al.*, 2017), *CROSSalive* can be integrated as a constrain in thermodynamics-based approaches such as *RNAfold*, which will allow study structural differences of RNAs *in vivo* and *in vitro* (Lorenz *et al.*, 2016).

## ACKNOWLEDGEMENTS

The authors thank Andrea Cerase and Alessio Colantoni.

---

**Supplementary Figure 1.** Cross-validation between neural networks trained on strong signal 100’000 icSHAPE fragments (equal size groups). The accuracy is reported for the 5% highest confidence set. Each comparison is done against the other datasets.

**Supplementary Figure 2.** Pipeline summarizing the process of filtering for protein contributions for m6a+ data.

**Supplementary Figure 3.** Pipeline summarizing the process of filtering for protein contributions for m6a-data.

**Supplementary Figure 4.** Distribution of *in vivo* minus *in vitro* icSHAPE scores along the entire murine transcriptome. Several regions are highly-variable, showing drastic changes from double-to single-stranded *in vivo* (score>0.5; d->s), or from single-to double-stranded (score<-0.5; s->d).

