## supplementary figures for "*CROSSalive*: a web server for predicting the *in vivo* structure of RNA molecules" (from another version of this preprint: version 1, 2019-05-05)

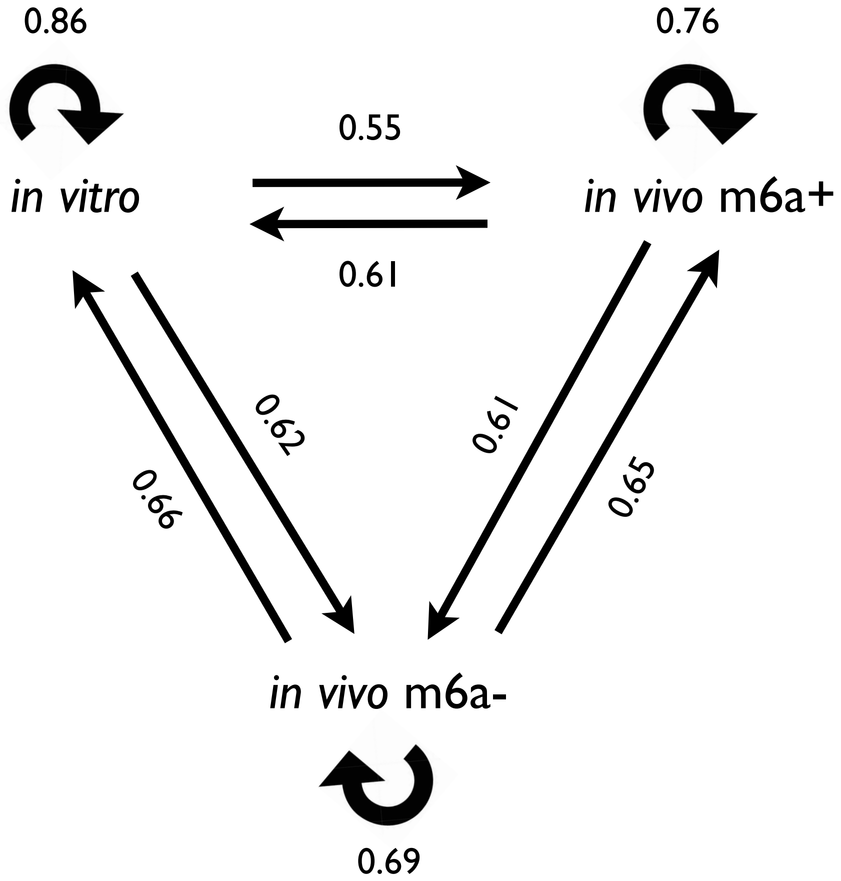

0.86
0.76
0.55
*in vitro*
*in vivo* m6a+
0.61
0.62
0.61
0.66
0.65
*in vivo* m6a-
0.69

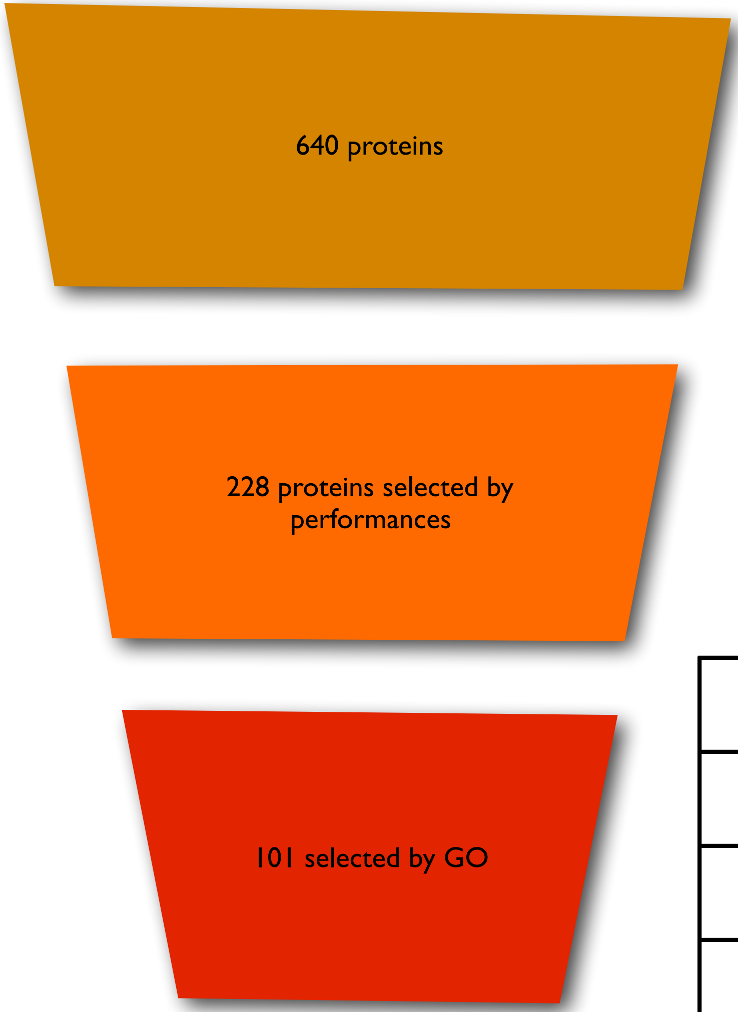


catRAPID

| Accuracy |
| --- |
| > 0.6 |

| GO identifier | GO name | p-value |
| --- | --- | --- |
| GO:0003725 | double-stranded RNA binding | 3.2x10-8 |
| GO:0003727 | single-stranded RNA binding | 2.7x10-7 |
| GO:0004386 | helicase activity | 7.8x10-8 |

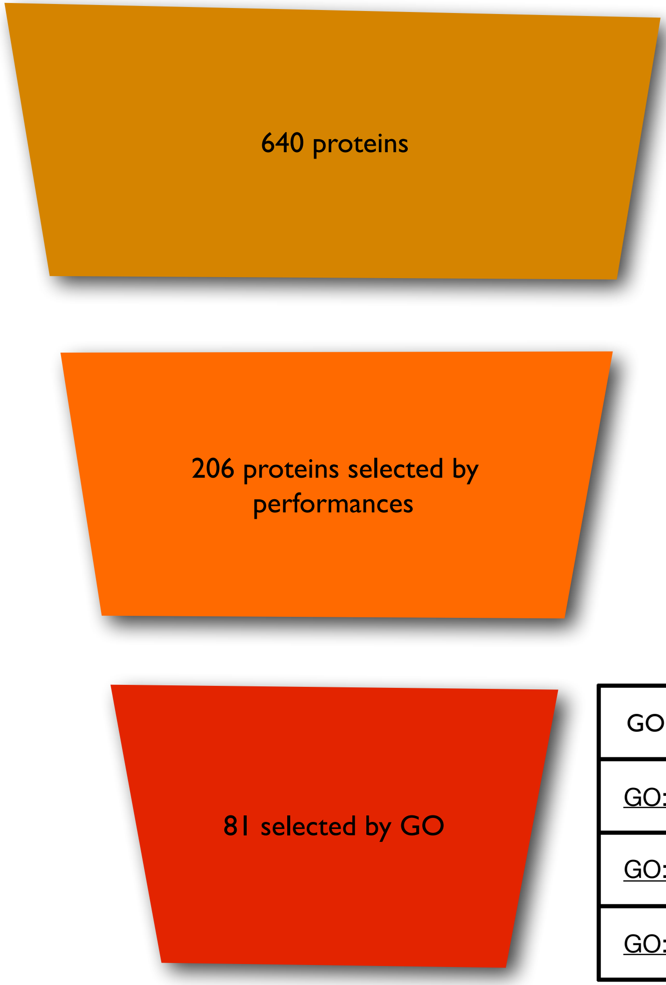


| Accuracy |
| --- |
| > 0.6 |

| GO identifier | GO name | p-value |
| --- | --- | --- |
| GO:0003725 | double-stranded RNA binding | 3.2x10-8 |
| GO:0003727 | single-stranded RNA binding | 2.7x10-7 |
| GO:0004386 | helicase activity | 7.8x10-8 |

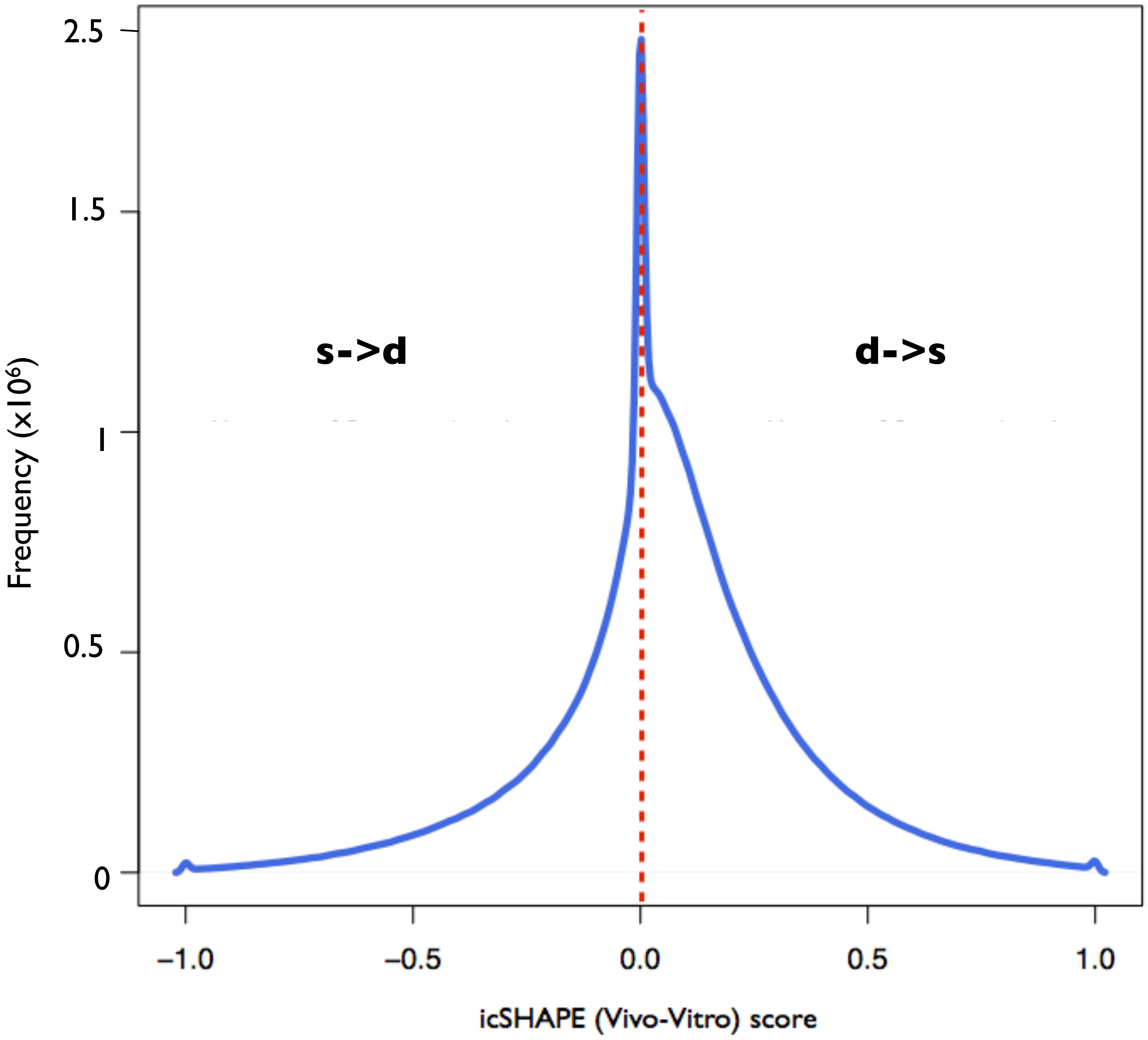

2.5
1.5
1
0.5
0
Frequency (×10⁶)
s->d
d->s
−1.0
−0.5
0.0
0.5
1.0
icSHAPE (Vivo-Vitro) score
